# Development of a semi-automated method for tumor budding assessment in colorectal cancer and comparison with manual methods

**DOI:** 10.1101/2021.06.17.448482

**Authors:** Natalie C Fisher, Maurice B Loughrey, Helen G Coleman, Melvin D Gelbard, Peter Bankhead, Philip D Dunne

## Abstract

Tumor budding is an established prognostic feature in multiple cancers but routine assessment has not yet been incorporated into clinical pathology practice. Recent efforts to standardize and automate assessment have shifted away from haematoxylin and eosin (H&E)-stained images towards cytokeratin (CK) immunohistochemistry. In this study, we compare established manual H&E and cytokeratin budding assessment methods with a new, semi-automated approach built within the QuPath open-source software. We applied our method to tissue cores from the advancing tumor edge in a cohort of stage II/III colon cancers (n=186).

The total number of buds detected by each method, over the 186 TMA cores, were as follows; manual H&E (n=503), manual CK (n=2290) and semi-automated (n=5138). More than four times the number of buds were detected using CK compared to H&E. A total of 1734 individual buds were identified both using manual assessment and semi-automated detection on CK images, representing 75.7% of the total buds identified manually (n=2290) and 33.7% of the total buds detected using our proposed semi-automated method (n=5138). Higher bud scores by the semi-automated method were due to any discrete area of CK immunopositivity within an accepted area range being identified as a bud, regardless of shape or crispness of definition, and to inclusion of tumor cell clusters within glandular lumina (“luminal pseudobuds”). Although absolute numbers differed, semi-automated and manual bud counts were strongly correlated across cores (ρ=0.81, p<0.0001). Despite the random, rather than “hotspot”, nature of tumor core sampling, all methods of budding assessment demonstrated poorer survival associated with higher budding scores.

In conclusion, we present a new QuPath-based approach to tumor budding assessment, which compares favorably to current established methods and offers a freely-available, rapid and transparent tool that is also applicable to whole slide images.

## Introduction

Tumor budding (TB) is the histological manifestation of local tumor cell dissemination, usually most evident at the invasive front region of a tumor mass. TB is an established prognostic factor in a number of solid tumors (1), although it has been most extensively studied in colorectal cancer (CRC). In pT1 CRC, the presence and extent of TB is predictive for nodal metastatic disease, and thus can be used as a clinical tool for identifying patients most likely to benefit from surgical resection (2). TB has also been shown to have prognostic value in all other stages of CRC, with most evidence reported for stage II disease (1,3,4).

Despite the potential clinical utility of TB, inconsistent qualitative criteria, definitions and non-standardized reporting have proven an obstacle to routine implementation in pathology practice and TB generally remains a “non-core” item in CRC reporting datasets (5–7). In an attempt to address this issue in 2016, the International Tumor Budding Consensus Conference (ITBCC) established a consensus definition of a tumor bud, namely a single tumor cell or tumor cell cluster of up to four cells, and an agreed histopathological method of assessment (8). Although encouraging data was emerging at that time regarding TB assessment by cytokeratin (CK) immunohistochemistry (IHC), most of the established evidence was based on haematoxylin and eosin (H&E) assessment. The consensus preference from ITBCC was for H&E staining in conjunction with a three-tier scoring system within a “hot spot” field area normalized to 0.785 mm^2^.

Since emergence of the consensus budding definition from ITBCC, there has been increased focus on standardization, reproducibility and automation, with a view to clinical implementation. This was the subject of a recent comprehensive review, which summarized twelve publications describing differing semi-automated approaches to TB assessment, almost all applied to CRC (9). Most used commercially-available software but two utilized open-source software (ImageJ), and some used a form of machine learning. Importantly, almost all were applied to CK IHC images, with only one method proposed for H&E. Other groups pursuing manual rather than semi-automated assessment of TB have also advocated for a CK IHC-based approach (10). However, a recent expert Delphi consensus process addressing TB concluded that more evidence was required before incorporating IHC into TB scoring (11).

One advantage of CK IHC over H&E assessment is the potential for greater reproducibility in overall TB grade (12), addressing a limiting step in progressing TB towards clinical implementation. While most studies have compared only overall TB grade, very few studies have examined TB assessment at the individual bud level, which is likely where most discordance lies. Recently, Bokhorst *et al* compared evaluation by a panel of seven ITBCC experts of 3000 candidate buds from CK-stained sections representing 46 patients with CRC and found only moderate agreement (13). Consensus classification was not reached on 41% of the candidate buds. Agreement was slightly better in this study for H&E assessment of individual buds compared with CK IHC, but far fewer H&E candidate buds were presented for evaluation.

In the current study, we compare manual H&E and CK assessment methods with a new semi-automated approach to TB assessment performed on digital images from a cohort of stage II and III colon cancers. Manual and semi-automated annotation of individual candidate buds on the same CK IHC images allowed scrutiny of discordance at the individual bud level and consideration of the optimal definition of a tumor bud for these methods of assessment. Results were analyzed for all methods against impact on survival, as a measure of relative performance and comparison of potential clinical utility.

## Materials and methods

### STUDY COHORT

The study utilized an established Northern Ireland population-based resource of n=661 stage II and III colon cancers, creation of which has been fully described previously (Northern Ireland Biobank ethical approval references NIB13-0069/87/88 and NIB20-0334) (14). The resource includes tissue microarrays (TMA), generated from representative tumor blocks containing the tumor advancing edge, with one 1 mm diameter core per tumor taken from a random area along the advancing edge. Although this does not reflect clinical practice, where TB grade is based on the “hotspot” area from within a representative whole tumor section, use of TMAs in this study allowed high throughput and representation of the full morphological spectrum of colon cancer.

3 µm sections from each TMA were stained with H&E and with an anti-cytokeratin immunohistochemical antibody (Cam5.2; Ventana, mouse monoclonal, Cell Conditioning 1 for 8 min, DAB chromogen) on a BenchMark ULTRA (Ventana Medical Systems Inc.) automated slide stainer. Glass slides were scanned on an Aperio AT2 Scanner (Leica Biosystems, Newcastle, United Kingdom) at x40 and imported into the open-source software QuPath (v0.2.3) (15) for evaluation. The scanned TMA images are available from the Northern Ireland Biobank (16) upon application.

The suitability of individual CK IHC-stained cores for inclusion was determined by manual visual assessment of the scanned images, after application of the QuPath TMA dearraying tool. Of note, TMA sampling from the advancing tumor edge is likely to generate a significant number of “misses”, with only peritumoral tissue sampled. Of the n=486 cores with sufficient tumor present and matched clinicopathological data, individual cores were also excluded if (a) only mucinous or signet ring cell carcinoma was present (n=26), (b) there were large areas of tumor necrosis (n=26) (c) tumor present exhibited weak, patchy or negative immunostaining (n=82), (d) there was significant stromal CK immunopositivity (n=25), or (e) tissue folding, fragmentation or any other technical artifacts precluded assessment (n=72) (Supplementary Figure 1). After the above exclusions, 255 cores remained for CK IHC evaluation. Manual H&E assessment for inclusion was performed after CK IHC assessment, and a further 61 cores were excluded, due to either a lack of tumor or tissue artifacts as described above, precluding H&E assessment. A further eight cases with less than one month of follow-up time were also excluded from the analysis. This left n=186 cases for analysis, having comparative TB data for all four methods of assessment, as detailed below, and clinicopathological data available including sufficient follow-up.

### MANUAL BUDDING ASSESSMENT

Buds were manually assessed on H&E and CK IHC images by an expert gastrointestinal pathologist (MBL). This process is depicted in Figure 1A-1E. Within QuPath, after dearraying, individual cores were shrunk by 30 µm to correlate with semi-automated assessment in excluding candidate buds touching the periphery of the core. Each individual bud was manually marked on all images using the point tool within QuPath, enabling quick and accurate quantification per core and the ability to review each individual bud counted. The ITBCC recommendations for H&E TB assessment were followed, with the only exception being that the TMA cores did not represent the budding “hot spots” for each tumor. However, each 1 mm diameter core approximates the ITBCC recommended 0.785 mm^2^ area for TB assessment (8). Furthermore, by using random cores from the advancing edge our analyses were tested in a wide range of morphological conditions. Pre-determination of the tumor region for assessment with the TMA approach allowed inter-method comparison of individual buds. “Pseudobuds” within areas of heavy acute inflammation were excluded as recommended (8,11).

**Figure 1.**
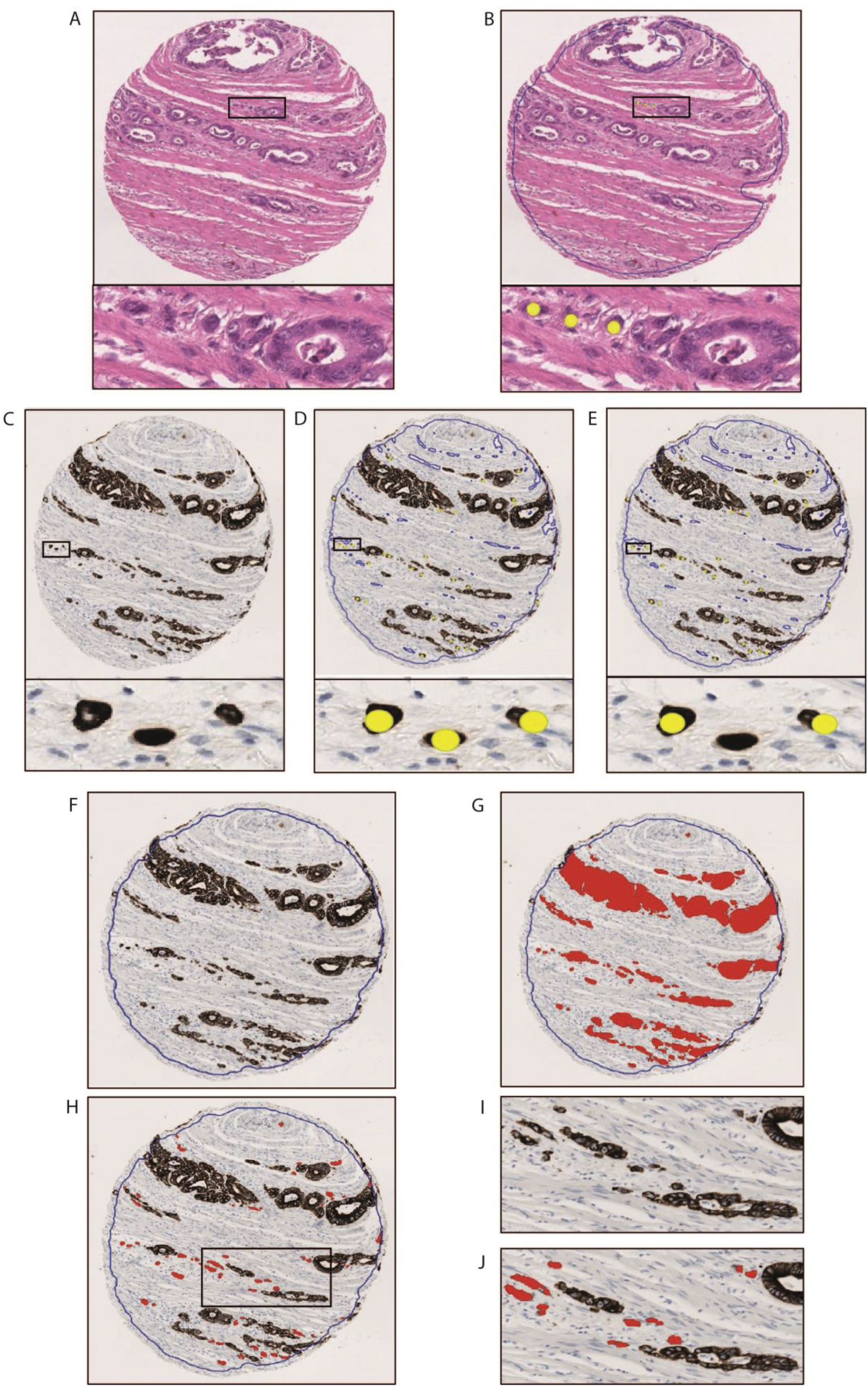
Methods of assessment of tumor budding. A-E, manual methods. F-J, semi-automated method. Tumor budding was manually scored on H&E-stained (A, B) and CK-stained (C-J) tissue microarrays. After dearraying, in all cores the tissue border was shrunk by 30µm to exclude candidate buds touching the periphery of the cores. Buds were annotated manually within QuPath (yellow dots in B, D and E). Initial CK buds (CK all, D) were revisited to exclude those lacking a region of nuclear pallor (CK pallor, E) and generate a second dataset applying this criterion. A semi-automated workflow was developed in QuPath (F-J). A binary classifier identified discrete CK positive regions (red). Lumens encapsulated by positive staining were filled in to exclude “luminal pseudobuds” (G). Buds were defined based on area of CK immunopositivity, the acceptable range (40-700 µm2) derived from analysis of the range of areas of the manually annotated CK buds. Objects with areas outside this range were excluded, leaving buds highlighted (H-J). (H&E, haematoxylin and eosin; CK, cytokeratin)

For initial manual assessment of CK-stained cores, the aim was to annotate as buds clusters of up to four tumor cells, as on H&E, accepting that visualizing and counting tumor cell nuclei is more difficult on CK IHC than on H&E (Figure 1C). Regions of irregular or ill-defined IHC staining were excluded, some considered likely to represent cellular fragments rather than viable buds. After this initial assessment was complete, annotated buds (CK all) were reassessed by the same observer to apply the recently suggested additional criterion of nuclear pallor in defining a bud (13). Those single cells or clusters lacking an identifiable region of nuclear pallor were removed to generate an additional budding dataset (CK pallor) which excluded objects lacking this potentially important feature (Figure 1E).

### SEMI-AUTOMATED BUDDING ASSESSMENT

The semi-automated method was based on a binary (immunopositive/immunonegative) threshold classifier built within QuPath (v0.2.3) and applied to the CK IHC images to identify tumor epithelium. This process is depicted in Figure 1F-1J. As before, following dearraying, individual cores were shrunk by 30 µm to exclude candidate buds touching the periphery of the core. All lumens completely encapsulated by positive staining were filled in, to prevent the detection of luminal tumor cells or cellular fragments mimicking buds (“luminal pseudobuds”) (Figure 1G). Color deconvolution was applied within QuPath to separate stains (17), followed by smoothing with a Gaussian filter and the application of a fixed global threshold to the deconvolved CK channel to identify connective discrete areas of immunopositivity (Resolution: 1.86µm/px; Channel: DAB; Prefilter: Gaussian; Smoothing sigma: 1.0; Threshold: 0.4). Buds were defined using this method not by number of tumor nuclei, but by area of CK immunopositivity. An acceptable range of bud area was derived from analysis of the range of areas of the manually annotated CK buds (described in detail below). Those objects with areas outside this range were excluded as buds (Figure 1H-1J).

### STATISTICAL ANALYSIS

Cox PH was conducted in Stata version 16 (Timberlake Consultants, StataCorp, College Station, TX, USA). All other analysis was conducted in R 4.0.3 (R Foundation for Statistical Computing, Vienna, Austria) (18). Statistical differences between the clinicopathological characteristics of the subset of patients utilized in this study compared to the overall cohort were determined. The Wilcoxon rank-sum test was applied to those groups with two levels, and Pearson’s Chi-squared test without continuity correction or Fisher’s exact test was applied to categorical variables where appropriate. The Kruskal-Wallis rank-sum test was used for the continuous variable.

Descriptive statistics were performed on the number of tumor buds detected per tissue core by each of the scoring methods. Spearman’s correlation coefficient was used to determine the strength of the linear relationship between each of the scoring methods. Univariable and multivariable analyses using the Cox proportional hazards regression model were performed to calculate hazard ratios (HR) and 95% confidence intervals (CI) for overall survival according to TB. Multivariable adjustments were age (<50, 50-<60, 60-<70, 70-<80, ≥80 years), sex (male, female), adjuvant chemotherapy receipt (yes, no), stage (II,III) and ECOG performance status (0-1, 2, 3-4, unknown). As the TMA cores in this study represent random cores from the tumor advancing edge, rather than TB hotspots, the ITBCC three category cut-offs are not strictly applicable. Therefore, survival analysis was conducted in two ways: (i) based on continuous bud counts to maximise statistical power, with per increment increases for each method based on relative ratios of total bud counts between methods; and (ii) applying modified ITBCC cut-offs to mimic categorization of scores for clinical decision making, and to generate Kaplan-Meier curves of prognostication, censored at five years of follow-up. ITBCC three category cut-offs were utilized for H&E scores (≤4, 5-9, ≥10 buds) and cut-offs for the other methods scaled up according to the TB score distribution for each method.

## Results

Of the original cohort, 186 individual cases were included in the study analysis. The overall clinicopathological characteristics are summarized in Table 1, which demonstrates that the subset of patient samples used in this current study shows no meaningful differences when compared to the overall stage II/III population-based cohort and can be considered a representative subset for analysis.

**Table 1.** Clinicopathological characteristics for the entire patient cohort and for the subset for study analysis. (ECOG, Eastern Cooperative Oncology Group; MSI, microsatellite instability; CRC, colorectal cancer; IR, interquartile range)

| Characteristic (n=entire/subset) | Entire cohort (n (%)) | Subset for analysis (n (%)) | p-value |
| --- | --- | --- | --- |
| **Sex (n=661/186)** |  |  | 0.26 |
| Male | 358 (54.2) | 92 (49.5) |  |
| Female | 303 (45.8) | 94 (50.5) |  |
| **Age, years (n=661/186)** | 73 (IR 64-79) | 74 (IR 65-80) | 0.18 |
| **ECOG performance status (n=410/113)** |  |  | 0.86 |
| 0-1 | 338 (82.4) | 92 (81.4) |  |
| 2 | 42 (10.2) | 11 (9.7) |  |
| 3-4 | 30 (7.3) | 10 (8.8) |  |
| **Tumor stage (n=661/186)** |  |  | 0.37 |
| II | 394 (59.6) | 104 (55.9) |  |
| III | 267 (40.4) | 82 (44.1) |  |
| **Tumor location (n=661/186)** |  |  | 0.42 |
| Proximal | 375 (56.7) | 116 (62.4) |  |
| Distal | 280 (42.4) | 69 (37.1) |  |
| Colon unspecified | 6 (0.9) | 1 (0.5) |  |
| **Tumor differentiation (n=657/183)** |  |  | 0.33 |
| Poor | 90 (13.7) | 20 (10.9) |  |
| Well/moderate | 567 (86.3) | 163 (89.1) |  |
| **Extramural venous invasion (n=610/163)** |  |  | 0.99 |
| Yes | 165 (27.0) | 44 (27.0) |  |
| No | 445 (73.0) | 119 (73.0) |  |
| **Microsatellite instability status (n=593/175)** |  |  | 0.30 |
| MSI-High | 136 (22.9) | 47 (26.9) |  |
| MSI-Low | 10 (1.7) | 5 (2.7) |  |
| Microsatellite stable | 447 (75.4) | 123 (70.3) |  |
| **Adjuvant chemotherapy (n=661/186)** |  |  | 0.85 |
| Yes | 186 (28.1) | 51 (27.4) |  |
| No | 475 (71.9) | 135 (72.6) |  |
| **Overall survival (n=661/186)** |  |  | 0.64 |
| Alive | 354 (53.6) | 96 (51.6) |  |
| Dead | 307 (46.4) | 90 (48.4) |  |
| **CRC specific survival (n=566/156)** |  |  | 0.82 |
| Alive | 354 (62.5) | 96 (61.5) |  |
| Dead | 212 (37.5) | 60 (38.5) |  |

### Deriving bud area range for semi-automated method

Semi-automated bud counts first required definition of an acceptable range of bud area, derived from analysis of the range of areas of the manually annotated CK buds. The semi-automated method initially identified all discrete areas of CK immunopositivity. Immunopositive areas, representing candidate buds, were initially captured over a wide size range (5-3000 µm^2^). Extremely small areas represented either tiny immunopositive tumor fragments, often in the context of gland rupture, (Figure 2A&2B) or non-specific immunostaining of uncertain nature (Figure 2C&2D). Large tumor areas were also annotated. By mapping the manual CK annotations to the semi-automated annotations, the areas of all manually annotated CK buds (CK all) could be measured within QuPath (Figure 2E&2F) and exported for analysis. The median CK bud area of the manually annotated CK buds (CK all), as measured by QuPath, was 225 μm^2^ (Figure 3A; interquartile range 133-388 μm^2^). The images, including manual and semi-automated annotations, of outliers at the low and high end of the area scale were reviewed, to explain implausibly small and large areas for some manually annotated buds. In some single cell buds, the semi-automated method excluded from the area measurement a prominent region of central nuclear pallor, thereby underestimating the true bud area (Figure 2G&2H). For some closely approximated buds, QuPath failed to resolve these as separate buds and considered their total combined area as a single immunopositive region, resulting in an apparent manually detected bud with a large area (Figure 2I&2J). Taking these erroneous extreme values into consideration, a range of 40-700 μm^2^ was chosen as acceptable in this study for defining a bud based on area of CK immunopositivity. Applying this definition, Figure 3B demonstrates by histogram the resultant areas and frequencies of the buds detected by the semi-automated method, having a lower modal bud area compared to the manual CK (CK all) method.

**Figure 2.**
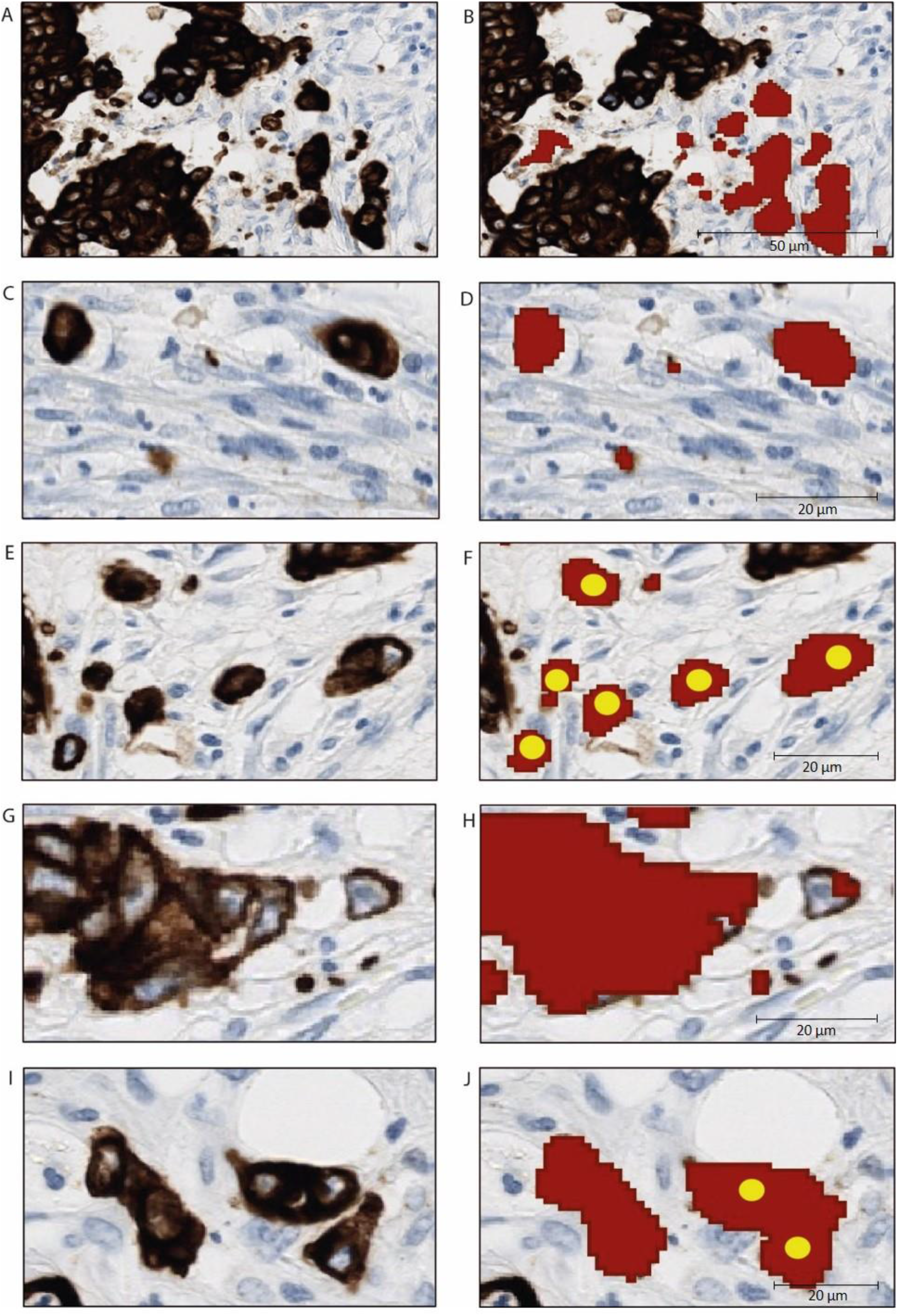
Deriving bud area range for semi-automated method. A, C, E, G and I: Unannotated images. B, D, F, H and J: Corresponding annotated images (red = QuPath annotations of CK) immunopositivity within broad area range of 5-3000 µm^2^; yellow = manual bud annotations). A and B: Tumor gland rupture, generating multiple tiny immunopositive fragments; C and D: Tiny immunopositive fragments of uncertain nature (arrows) detected alongside two true tumor buds (arrowheads); E and F: Six manually annotated (CK all) tumor buds, with areas measured by QuPath (range 107-384 µm^2^). G and H: A manually annotated single tumor cell bud with prominent nuclear pallor resulting in underestimation of the bud area by QuPath (measured as 10 µm^2^). I and J: Two closely adjacent buds annotated manually, but considered by QuPath as one large immunopositive area (measured as 820 µm^2^). (CK, cytokeratin)

**Figure 3.**
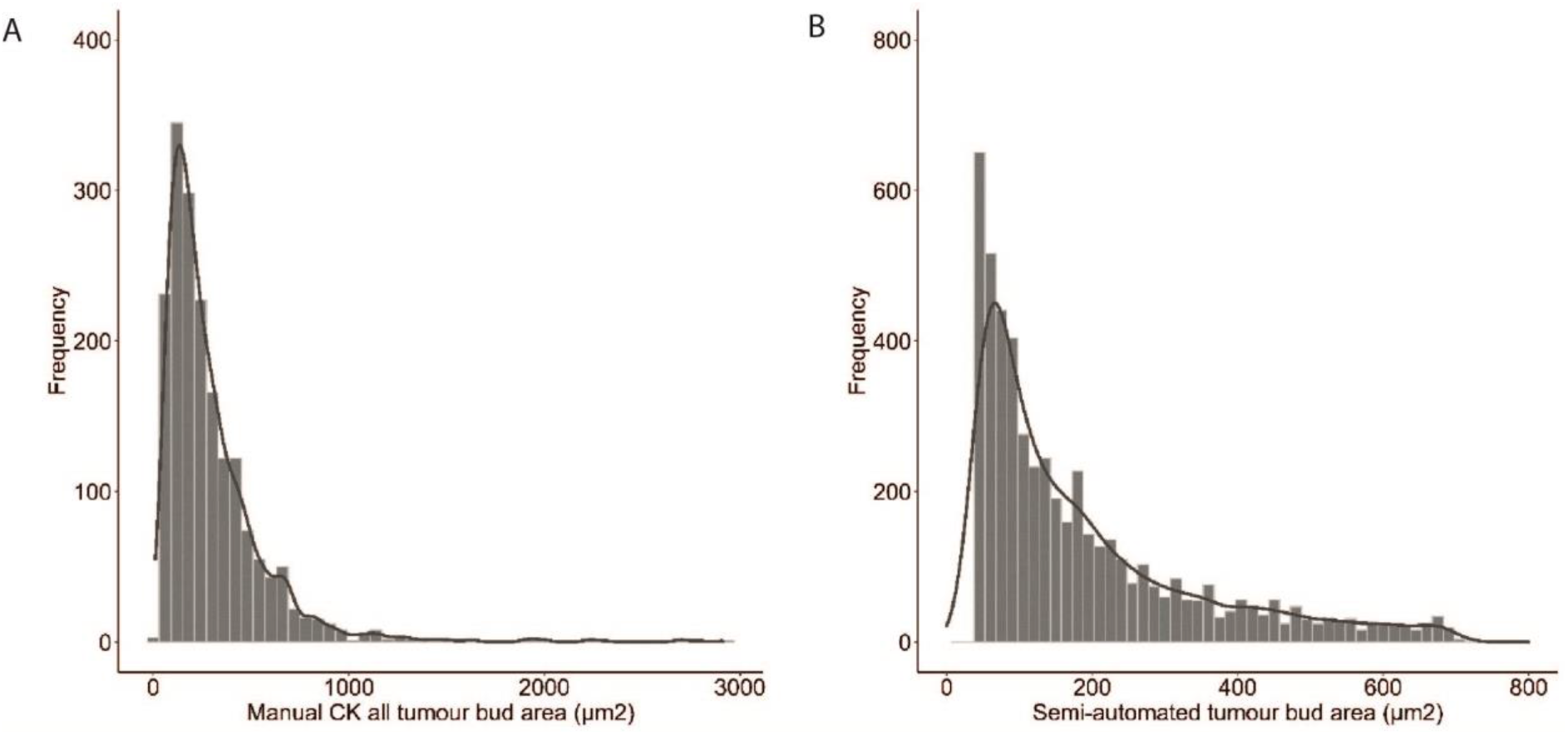
A. Histogram of manual CK-detected bud areas, as measured by QuPath. This was the basis of selecting a suitable area range to define a bud applying the semi-automated assessment method. B. Histogram of areas of buds detected by the semi-automated method, applying a range of 40-700 µm2. (CK, cytokeratin)

### Total bud count comparisons

The total number of buds detected by each method (Figure 4A), over the 186 TMA cores, were as follows; manual H&E (n=503), CK all (n=2290), CK pallor (n=1825) and semi-automated (n=5138). These findings indicate that more than four times the number buds were detected using CK (CK all) compared to H&E, and more than three times the number if restricting to those buds with central pallor (CK pallor). The semi-automated method detected over ten times more buds than H&E and over twice as many buds as CK (CK all). Comparing bud totals and frequencies for each method showed progressively increasing numbers of cases with higher numbers of buds moving from H&E to CK to semi-automated assessments (Figure 4B). Comparison of total bud numbers between H&E and CK showed moderate correlation (Figure 4C, ρ=0.60, p<0.0001), whereas strong correlation was observed between CK all and semi-automated methods (Figure 4D, ρ=0.81, p<0.0001).

**Figure 4.**
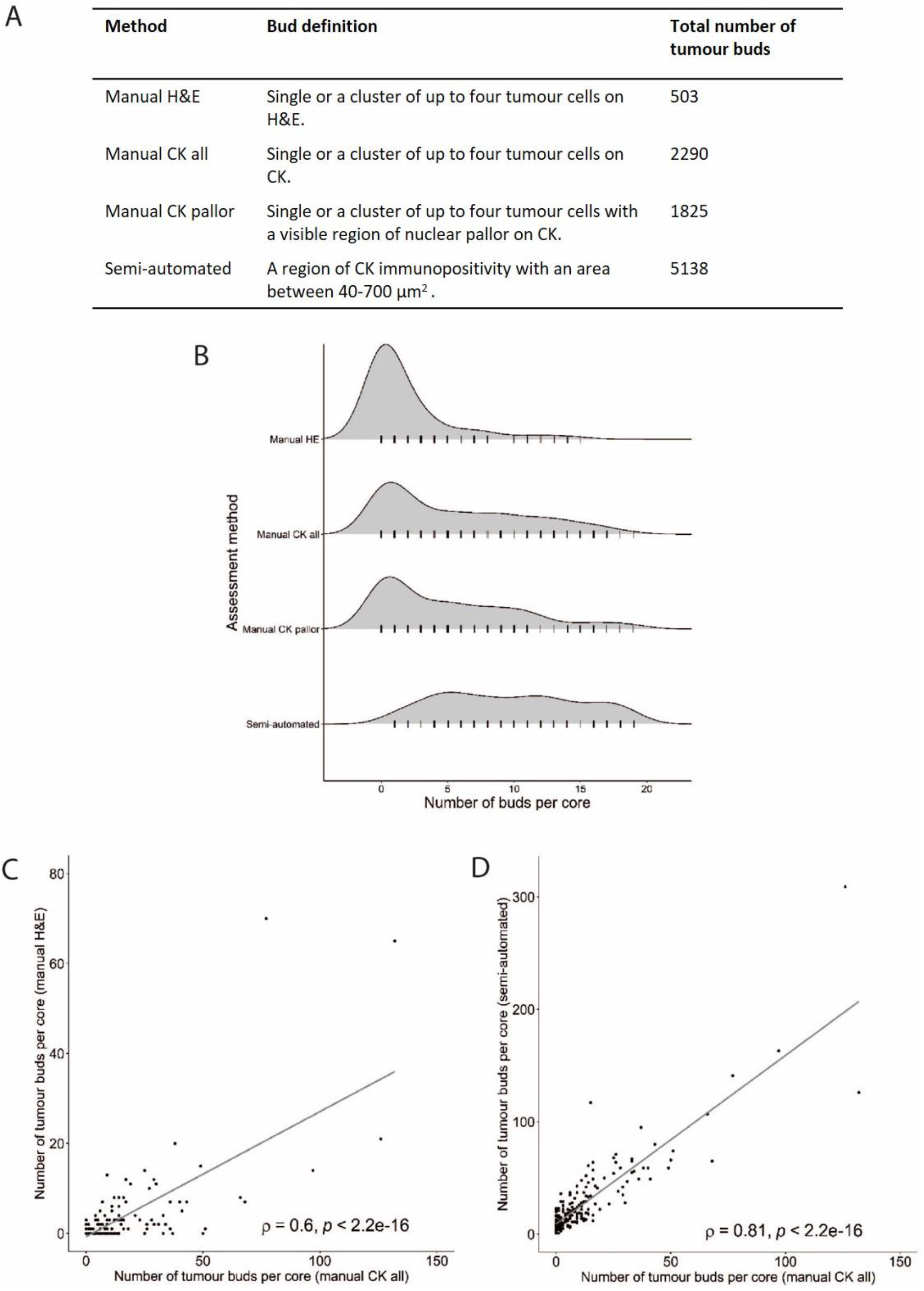
Total bud count comparison across all scoring methods. A. Total number of tumor buds detected by each of the four methods of assessing tumor budding in all study cases (n=186). B. Total buds per core and frequencies of each number within the study group, for cores with up to 20 buds detected. C. Correlation of total bud counts per core assessed manually by H&E and CK (CK all). D. Correlation of total bud counts per core assessed on CK manually (CK all) and by the semi-automated method. (H&E, haematoxylin and eosin; CK, cytokeratin)

### Bud by bud comparisons

As both manual CK assessments and the semi-automated assessment were performed on the same set of images, bud by bud comparison was possible for these methods. A total of 1734 individual buds were identified both by manual assessment (CK all) and semi-automated detection, representing 75.7% of the total manual buds identified (n=2290) and 33.7% of the total semi-automated buds detected (n=5138) (Figure 5). Accepting the manual CK method as the relevant gold standard, these equate to the sensitivity and positive predictive value respectively of the semi-automated method for detection of CK (CK all) buds.

**Figure 5.**
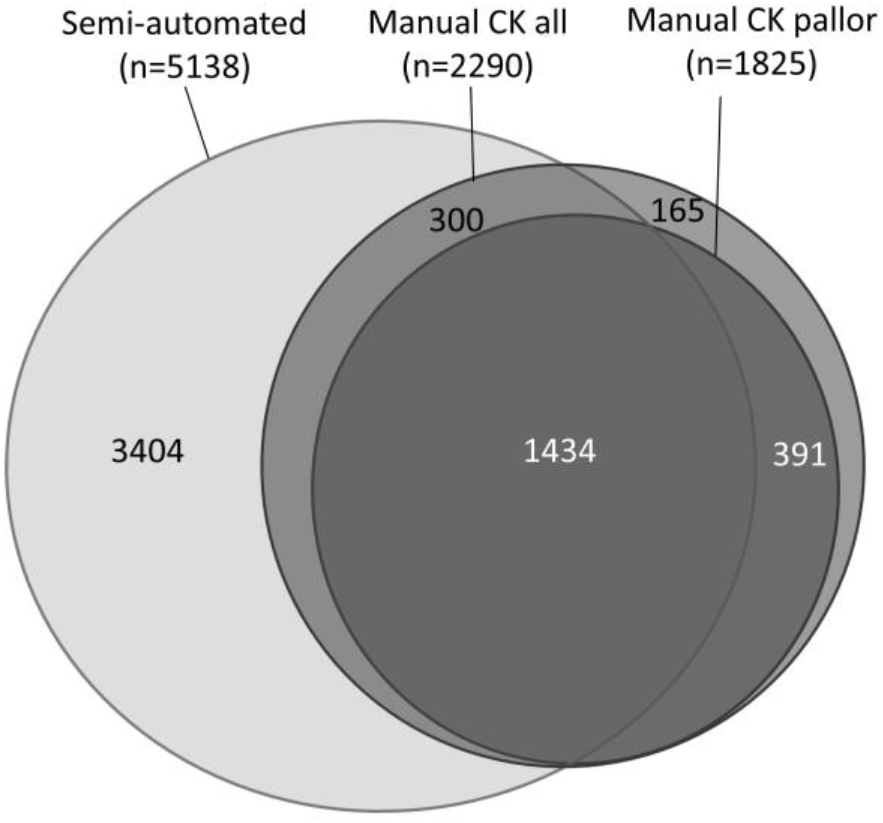
Individual bud by bud comparison of manual CK all, manual CK pallor and semi-automated assessment methods. (CK, cytokeratin)

### Bud discordance between methods

Many tumor areas demonstrated excellent concordance, with buds being detected by both manual CK and semi-automated assessment methods after application of the specified area range for the semi-automated method (Figure 6A&6B). However, elsewhere concordance between these assessment methods was poor. This was in large part due to the semi-automated method accepting as a bud any discrete area of CK immunopositivity within the accepted area range, regardless of shape or crispness of definition, features which would typically be considered in the manual assessment of a bud (Figure 6C&6D). The other main explanation for much greater numbers of buds by the semi-automated method relates to “luminal pseudobuds”. Manual assessment discounts as buds, tumor cells or clusters lying within glandular lumina. When surrounded by circumferential staining, QuPath was able to fill in the glandular lumina, to avoid counting such mimics as buds (Figures 1F&1G, 6E&6F). However, when staining was not circumferential, QuPath counted these luminal immunopositive fragments as buds (Figure 6G&6H). This was a particular problem at core peripheries, where the complete gland circumference was not captured within the core (Figure 6I&6J). The inclusion of the more stringent nuclear pallor criterion to define a CK bud by manual assessment had a minor additional impact on the discordance in bud numbers between manual CK and semi-automated assessments (Figure 5).

**Figure 6.**
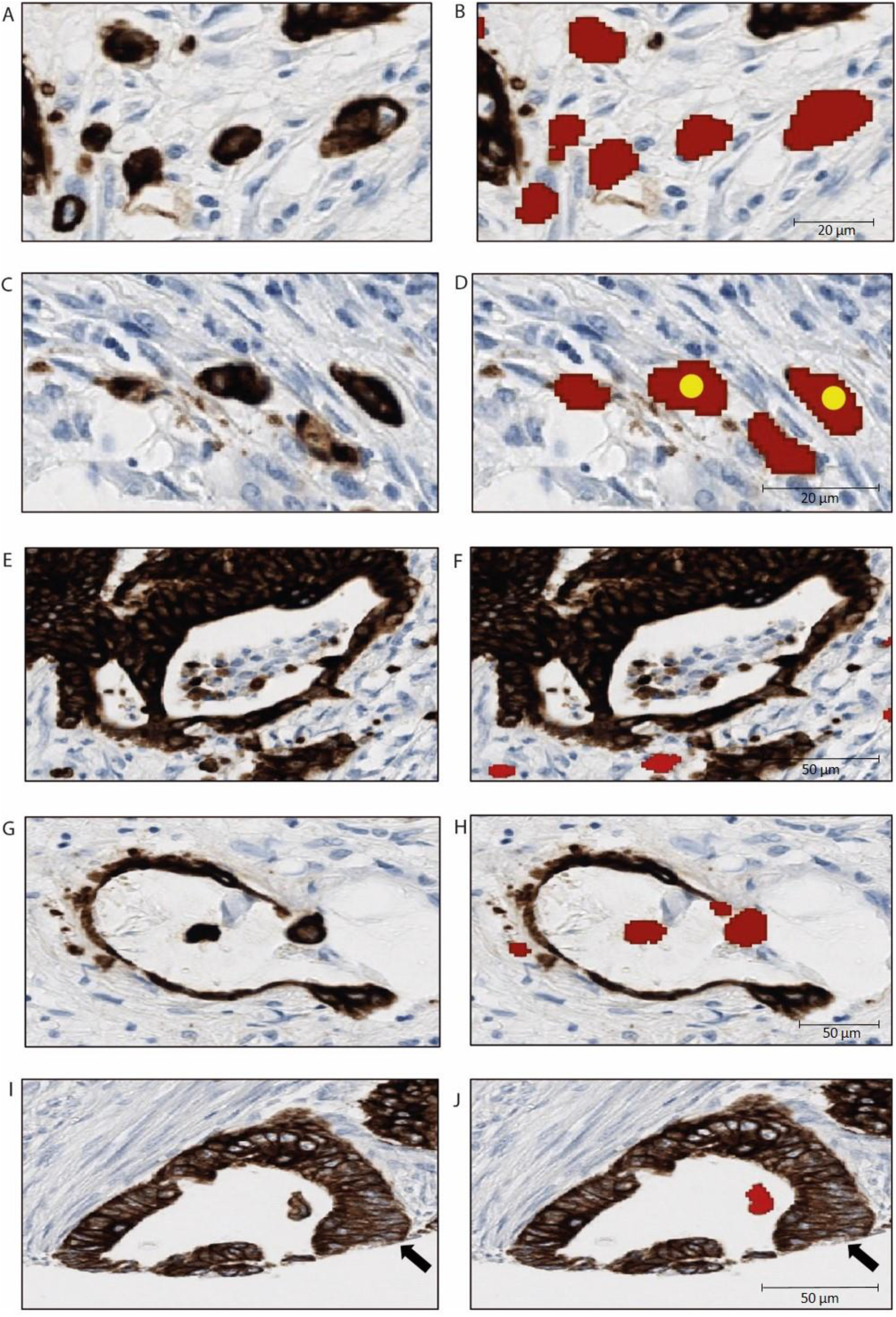
Discordance in bud assessment between manual CK and semi-automated methods. A, C, E, G & I: Unannotated images; B, D, F, H & J: Corresponding annotated images (red shapes = QuPath bud annotations; yellow circles = manual bud annotations). A & B: Perfect concordance in annotation of six tumor buds between manual (CK all) and semi-automated methods; C & D: Poor concordance, with the manual method identifying two buds and QuPath identifying two additional, less well-defined, buds; E & F: Mimics of tumor buds within complete glandular lumina (“luminal pseudobuds”) are discounted as buds by both manual and semi-automated methods, resulting in concordance; G & H, I & J: If glands are disrupted, resulting in incomplete circumferential immunostaining, QuPath cannot “fill in” the gland lumen and these luminal mimics are counted as buds by the semi-automated method, a particular problem in tissue microarrays when glands involve the core edge (arrow, in I & J). (CK, cytokeratin)

A smaller number of manual CK buds (CK all and CK pallor) were not detected by the semi-automated method. These are explained by erroneous bud area measurement, as described above. Incorrect assessment of true bud area, because of exclusion of a region of nuclear pallor (Figure 2G&2H) or failure to resolve closely adjacent buds (Figure 2I&2J), generated areas below or above the accepted range, and thereby failure to identify these manually detected buds by the semi-automated method.

### Survival analysis

Of the n=186 patients included in the analysis, by the end of follow-up (mean ± standard deviation, 5.5 ± 3.0 years; range 0.12-10 years), 90 had died of which 60 were from a CRC-related cause. All four methods of TB assessment demonstrated reduced survival associated with higher budding scores (Table 2). HRs were similar for both of the CK methods and for the semi-automated method in the univariable (manual CK all: HR 1.09, 95%CI 1.05-1.14; manual CK pallor: HR 1.11, 95%CI 1.06-1.18; semi-automated: HR 1.09, 95%CI 1.04-1.14) and multivariable (manual CK all: HR 1.06, 95%CI 1.02-1.11; manual CK pallor: HR 1.08, 95%CI 1.02-1.14; semi-automated: HR 1.06, 95%CI 1.01-1.11) models, and slightly lower for the H&E method in both univariable (HR 1.03, 95%CI 1.01-1.05) and multivariable (HR 1.02, 95%CI 1.00-1.04) models. All findings were statistically significant aside from H&E findings in the multivariable model.

**Table 2.** Univariable and multivariable Cox proportional hazards overall survival analysis comparing four methods of tumor budding assessment. (H&E, haematoxylin and eosin; CK, cytokeratin; CI, confidence intervals)

| Assessment method | Overall survival |  |
| --- | --- | --- |
|  | Hazard ratio (95%CI) |  |
|  | Unadjusted | Multivariable adjusted model |
| **Manual H&E** |  |  |
| Per 1 bud increase | 1.03 (1.01-1.05) | 1.02 (1.00-1.04) |
| *P-value* | *0.001* | *0.08* |
| **Manual CK all** |  |  |
| Per 5 bud increase | 1.09 (1.05-1.14) | 1.06 (1.02-1.11) |
| *P-value* | *<0.001* | *0.004* |
| **Manual CK pallor** |  |  |
| Per 5 bud increase | 1.11 (1.06-1.18) | 1.08 (1.02-1.14) |
| *P-value* | *<0.001* | *0.005* |
| **Semi-automated** |  |  |
| Per 10 bud increase | 1.09 (1.04-1.14) | 1.06 (1.01-1.11) |
| *P-value* | *<0.001* | *0.01* |

Kaplan-Meier survival analysis showed patients with higher TB grades had significantly reduced overall five year survival, when assessed by any of the four methods presented (Figure 7). Stratification was lesser for H&E assessment (p=0.026) than the other three methods, all of which were comparable (p<0.0001, P<0.0001, p=0.0009). Introduction of nuclear pallor to the manual CK assessment did not meaningfully impact stratification.

**Figure 7.**
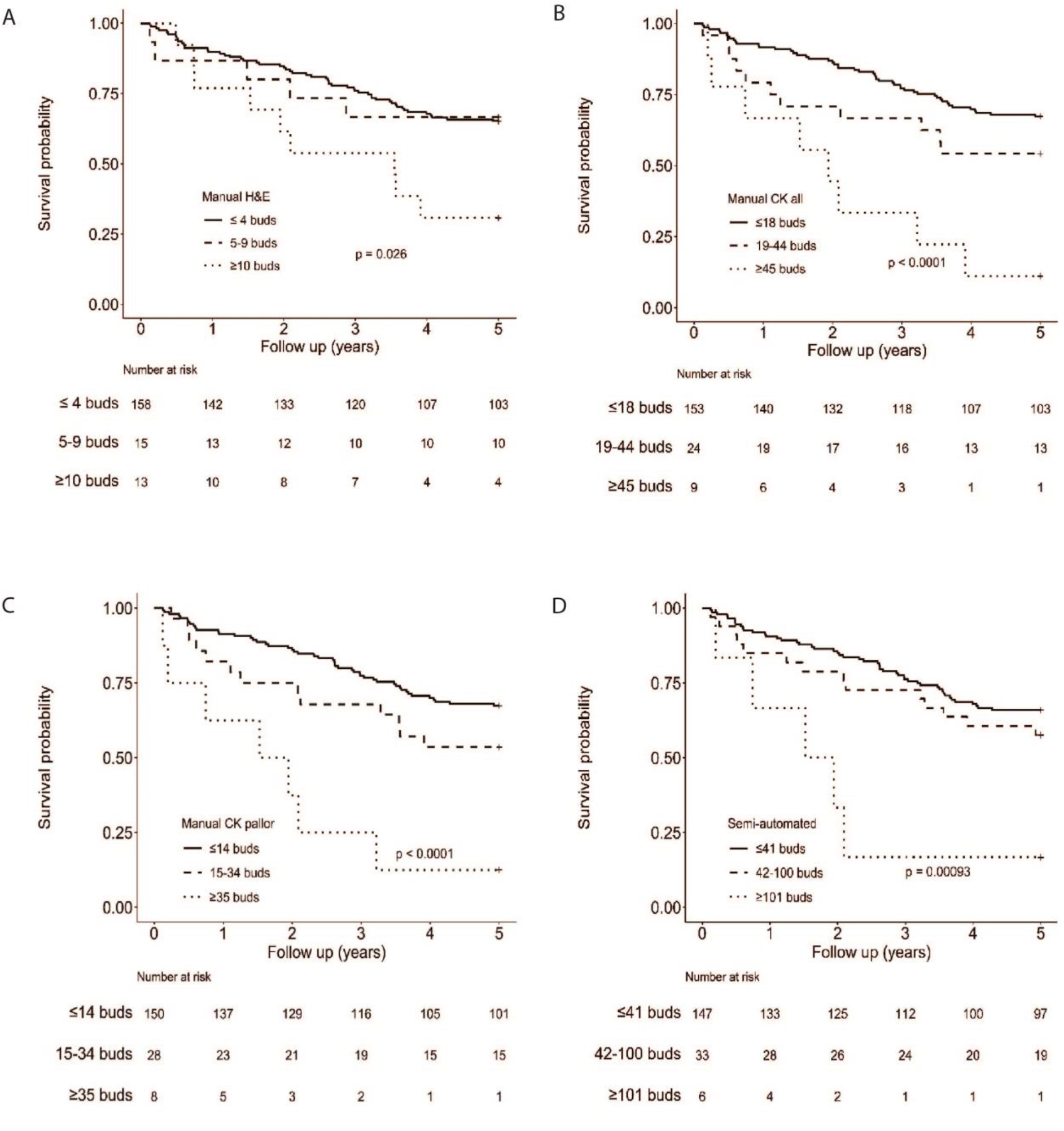
Kaplan-Meier estimates demonstrating overall survival differences in patients with stage II/III colon cancer according to low, moderate and high grade tumor budding assessed by four different methods, with recommended category cut-offs scaled according to budding score distribution for each method.

## Discussion

TB is well established as an adverse prognostic feature in CRC in several clinical settings (1). Despite considerable existing evidence in this regard, assessment of TB has not yet been incorporated into routine clinical practice. In large part, this is because of uncertainty regarding the most appropriate method of assessment, specifically the most appropriate stain for counting buds and whether to persist with manual assessment or adopt some form of semi-automated approach. In this study, we used QuPath to develop a new digital pathology-based semi-automated TB assessment tool for CK-stained sections, which we then compared to established methods of TB assessment in a cohort of colon cancers using a TMA approach. As the study included TMA cores from the tumor advancing edge of stage II/III colon cancers, rather than the budding hotspot advocated for clinical use, the primary focus of this paper was a bud by bud comparison of manual CK and our semi-automated assessment method, rather than to provide further evidence of adverse prognostic significance of TB.

Our data indicates that CK IHC detected over four times more buds than H&E-based assessment of parallel sections, which is consistent with previous studies observing three to six times more buds with CK IHC than with H&E staining (12). Although not examined in this study, it is postulated that CK IHC is particularly valuable in highlighting single cell buds and distinguishing these from epithelioid stromal or histiocytic cells by indicating their epithelial cell lineage, less readily apparent on H&E. Bokhorst *et al* have hypothesized that inter-observer variability on H&E assessment may be more problematic for single cell buds than for two to four cell buds (13). H&E assessment allows better evaluation of the microenvironment surrounding buds and so it is possible that a further reason contributing to fewer H&E buds relates to greater exclusion of so-called pseudobuds at sites of active inflammation, often related to gland rupture (1). The inflammatory environment is less readily appreciated in CK IHC preparations, meaning pseudobuds may be less identifiable and therefore less likely to be excluded.

The threshold semi-automated approach identified approximately 2.5 times more buds than manual CK assessment. Higher bud counts have been observed previously when comparing a semi-automated to manual CK assessment method, but without quantification (19). In data presented here, we find that bud by bud comparison revealed only moderate agreement between these two assessment methods for individual buds. Some of the discrepancy might be explained by the tendency of any human observer to err slightly on the side of under-counting, either through occasionally missing a possible true bud or by making a conservative judgement in an ambiguous case. By contrast, one can expect a threshold-based approach that defines a bud by area of CK immunopositivity to err definitively on the side of overestimation, consistently including more irregular or ill-defined ambiguous tumor cell clusters. It is possible that incorporating further criteria into the bud definition may improve agreement between semi-automated and manual assessments, such as a measure of circularity (20). However, given that there is no *a priori* reason to suppose buds are circular, this can introduce further subjectivity. In this study we have aimed to minimize the adjustable parameters, relying primarily upon a staining threshold and area filter to achieve a replicable baseline of quantitative assessment. The area range we selected to define a tumor bud (40-700 μm^2^) was based on the corresponding area range of manually detected CK buds, which is wider than that chosen by Takamatsu *et al.* (100-480μm^2^) but narrower than that chosen by Bokhorst *et al.* (25-1000μm^2^) (13,20). This already indicates the lack of accepted parameters in defining bud characteristics through image analysis, although such parameters will inevitably have a profound influence upon the absolute numbers of buds detected. Interestingly, we found that, despite the substantial differences in absolute bud counts between methods of assessment, correlation remained high – suggesting that the signal remains high amidst the noise.

As there is evidence to support high TB as an adverse prognostic factor across all stages of CRC (1,3,4), survival analysis was conducted applying the four methods of TB assessment, as a measure of comparative performance. Despite the limitations of random core sampling, TB assessed by all four methods was, as expected, significantly associated with reduced overall survival at five years of follow-up. This association was weakest for H&E assessment, and non-significant on the multivariable model, but it is likely that H&E assessment, with the lowest bud counts in general, will have been impacted more by the random core approach in our study in comparison to the other methods yielding much higher bud counts. Nevertheless, the other three methods all stratified patients better than H&E with respect to survival and achieved almost identical hazard ratios based on evaluation of continuous bud counts. Importantly, despite its simplicity and only moderate agreement with manual CK assessment for individual buds, the semi-automated threshold approach in QuPath provided an association between higher grades of TB and worse overall patient survival, even when applied to random tumor cores.

A recent modified Delphi process conducted amongst an international group of expert gastrointestinal pathologists supported ongoing assessment of TB using H&E-stained slides, with more evidence required to move to IHC, but also suggested that digital image analysis was likely to facilitate implementation into clinical practice (11). As almost all TB algorithms published to date rely on CK rather than H&E-stained images, it seems likely that the optimal approach will ultimately be one based on evaluation of the most representative tumor section, stained for CK. With increasing developments in digital pathology and growing access to digital whole slide images in routine practice, some form of semi-automated approach is attractive for reasons of efficiency, cost and reproducibility. Such semi-automated methods can be easily applied over a much larger tumor area to accurately identify the budding density over any agreed area denominator. The consensus 0.785 mm^2^ area applicable to microscopy is less relevant to whole slide image analysis. Nevertheless, most current evidence for TB significance is based on this hotspot area, and correlation with microscopy assessment of TB will be important for the foreseeable future.

It is likely that the semi-automated approach to budding assessment described in this study is overly simplistic for clinical use as it is unable to detect some of the more subtle morphological features of tumor buds, such as nuclear pallor, nor exclude mimics such as pseudobuds. Future clinical implementation will require more refined methodologies, likely involving deep learning (9,21), however as yet no such method is widely available to the TB community. The semi-automated QuPath approaches developed and applied in this study will be of potential benefit to ongoing translational TB research in retrospective cohorts as a much cheaper, more efficient and readily customizable open-source method compared to commercial software solutions. Such tools can be utilized either as a standalone TB assessment or as an adjunct to developing more sophisticated methods for example by identifying large numbers of candidate buds for consensus expert evaluation, classification and application to training of deep learning algorithms.

Assessment of TB by CK IHC has been shown by some studies to improve inter-observer reproducibility, an important requirement when considering incorporation of any new parameter into routine pathology practice (12,22). However, a recent study employing CK IHC for TB assessment examined inter-observer agreement at the individual bud level and found only moderate agreement, no better than for H&E assessment (13). The authors considered two reasons for this: firstly, that individual tumor nuclei within immunopositive clusters are sometimes difficult to discern, and therefore count, on CK IHC; and secondly, that the surrounding inflammatory environment is more difficult to assess on CK IHC than on H&E, making evaluation of potential “pseudobudding” more challenging. Less evidence is available on reproducibility of semi-automated methods but it is intuitive that more automation implies greater reproducibility. Takamatsu *et al.* found significantly better reproducibility amongst three pathologists with their semi-automated method (kappa coefficient = 0.781) compared to manual assessment (kappa coefficient = 0.463) (20). Nevertheless, some degree of manual oversight remains important whilst new methods are developed and tested.

Introducing the additional criterion of nuclear pallor into the manual CK assessment method made no meaningful alteration to the resultant hazard ratio (CK pallor HR 1.11, 95% CI 1.06-1.18; CK all HR 1.09, 95% CI 1.05-1.14) or the Kaplan-Meier survival stratification, providing no real evidence from this study for inclusion of this criterion. Previously suggested by Bokhorst *et al.* (13), to help exclude CK positive non-viable tumor cell fragments from consideration as buds, this feature should be the focus of future studies based on hotspot TB assessment on whole tumor sections from appropriate CRC cohorts, to ascertain the potential impact of this morphological criterion on clinical relevance of TB and inform future discussions on bud definition.

This study is limited by the random nature of the tumor core samples, limiting analysis of the clinical significance of TB scores with respect to survival analyses, and by the single pathologist manual assessment of buds without any ability to assess reproducibility. However, a detailed comparison of different TB assessment methods is described, applied to a wide morphological spectrum of colon cancers, with bud by bud comparison between methods.

Although our CK thresholding approach resembles methods applied in previous TB studies (9,20,23), to our knowledge the current study is the first to describe an interactive tool for TB assessment that is freely available, open-source, and can be readily applied to whole slide images as part of a full analysis workflow. This is possible because of the extensive additional functionality within QuPath, including the ability to precisely define regions of interest (e.g. a 1 mm boundary delineating the tumor advancing edge), identify hotspots, and export quantitative metrics. These features are illustrated in Figure 8, applying the methods adopted in this study to a whole slide image from a sample CRC case rich in tumor buds. Manually-derived and semi-automated budding density “heat maps” are almost identical. In contrast to assessment approaches driven entirely by machine learning, which can be confounded by even subtle variations in staining or scanning (24,25), our comparatively simple thresholding method can be readily adapted to new images by adjusting a small number of intuitive parameters – making it immediately accessible to any laboratory wishing to apply the technique. Nevertheless, it is clearly desirable to achieve a better discrimination of true buds from false positives. In this regard, QuPath’s generic support for machine learning, previously described for cell classification (15), can be incorporated into a more elaborate analysis workflow. Having established in this study the first open and replicable end-to-end analysis protocol for TB assessment suitable for whole slide images, we aim to collaborate with other groups to develop a refined, open-source bud identification algorithm based upon a more diverse training dataset across multiple centers.

**Figure 8.**
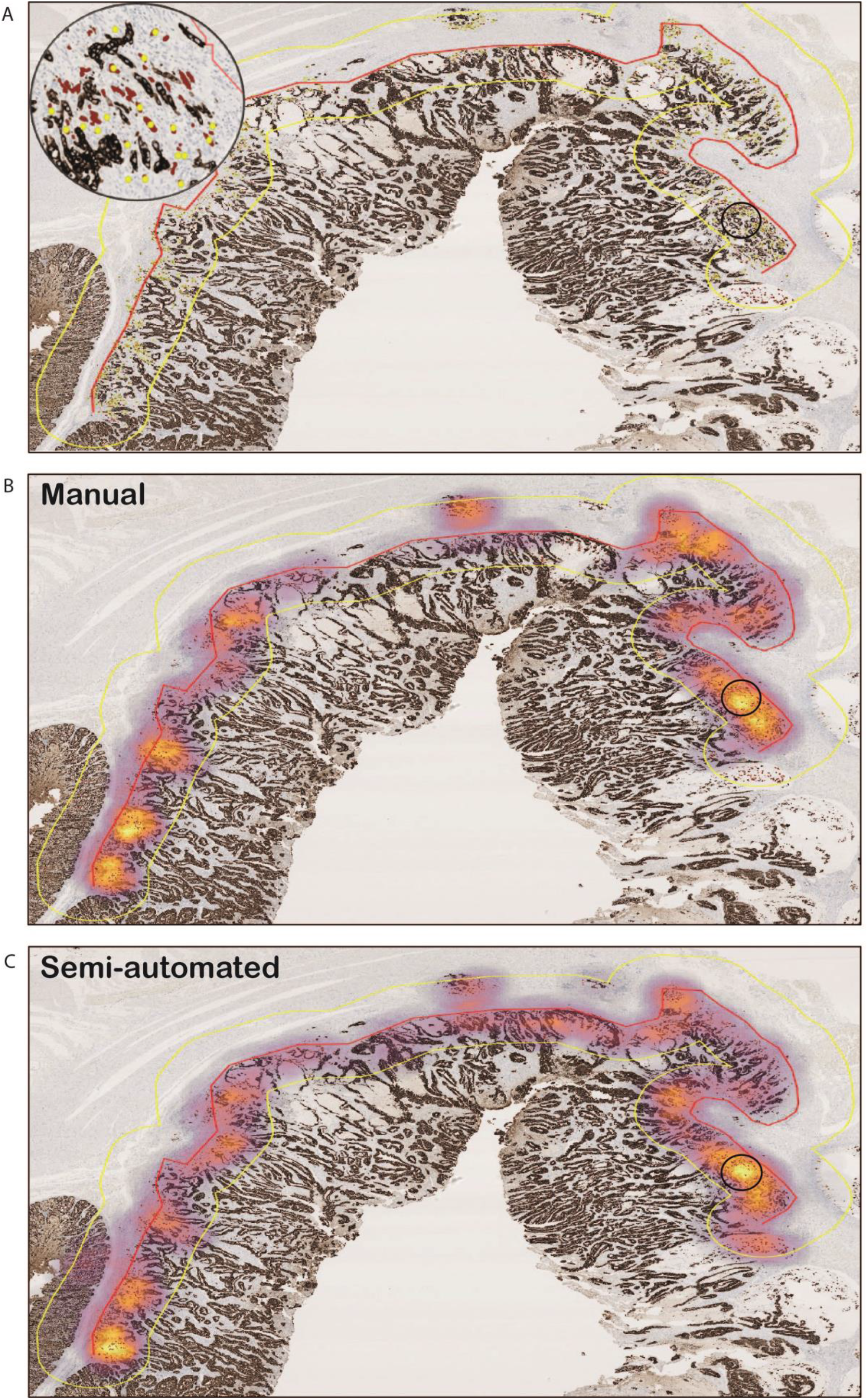
Tumor budding assessment applied to a whole slide CK-stained image of colorectal cancer. A high budding case has been chosen for illustration. A: after manual annotation of the advancing edge (red line) using the QuPath line tool, the expand annotation tool is used to expand the annotation 1 mm inwards and outwards, delineating the tumor advancing edge region of interest (yellow line) for budding assessment. Manually identified (yellow circles) and independently detected QuPath (red shapes) buds are shown (magnified in inset for “hotspot” area); B: bud density heat map based on manual bud annotations; C: bud density heat map based on QuPath bud annotations. Density colormaps are normalized independently for each image according to the maximum bud density within the image. The 0.785 mm^2^ “hotspot” is highlighted (black circle) in each image. (CK, cytokeratin)

In conclusion, we present a new QuPath-based approach to TB assessment. This demonstrates moderate agreement with manual CK-based assessment at a bud-by-bud level and comparable ability to stratify a cohort of patients with stage II/III colon cancer for overall survival. More importantly, it shows QuPath’s potential as a freely-available, rapid and transparent tool for TB assessment, applicable to whole slide images, which can be used in translational research as a standalone method or as an aid in developing future approaches suitable for clinical implementation.

## Acknowledgements

The samples used in this research were received from the Northern Ireland Biobank which has received funds from HSC Research and Development Division of the Public Health Agency in Northern Ireland and the Friends of the Cancer Centre.

The Belfast Health and Social Care Trust Department of Cellular Pathology is acknowledged for assisting with immunohistochemistry.

## Ethics Approval and Consent to Participate

Ethical approval for this study was granted by the Northern Ireland Biobank (NIB13-0069/87/88 and NIB20-0334).

## Author Contributions

Study concept and design: N.F., M.L., P.B. and P.D.; development of methodology N.F., M.L., P.B. and P.D.; QuPath software development: M.G. and P.B.; acquisition, analysis and interpretation of data and statistical analysis: N.F., M.L., H.C., and P.D; writing initial draft: N.F. and M.L.; review and revision of subsequent drafts, N.F., M.L., H.C., M.G., P.B. and P.D. All authors read and approved the final manuscript.

## Data Availability Statement

All codes used in this current study are publicly available at https://github.com/petebankhead/qupath-budding-scripts. The images and datasets used in the current study are available on application to the Northern Ireland Biobank.

