## Supplementary figures and images for "Development of a semi-automated method for tumor budding assessment in colorectal cancer and comparison with manual methods"

### Supplementary Figure 1

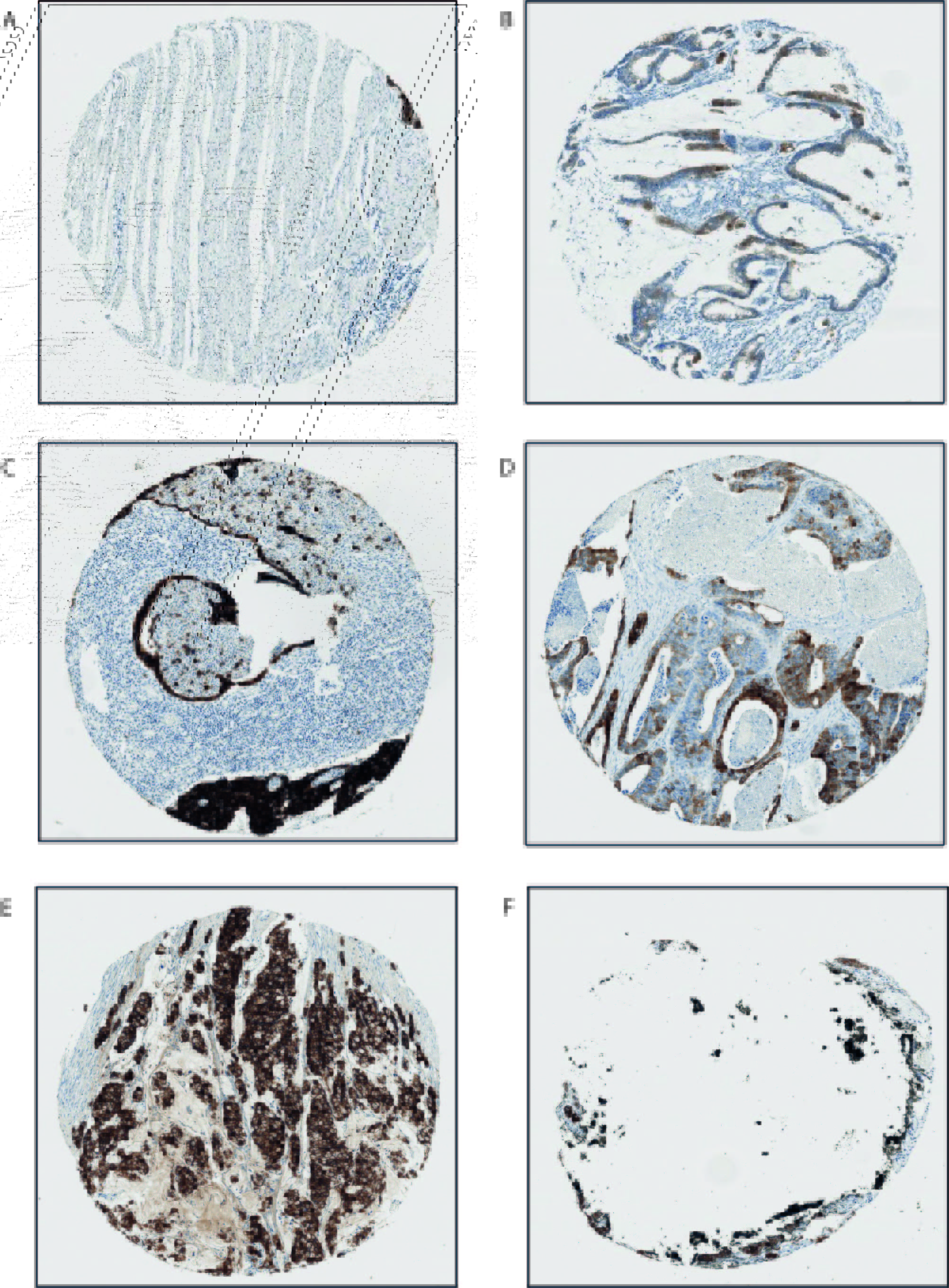
